# Annexin A2 depletion exacerbates the intracerebral microhemorrhage induced by acute rickettsia and Ebola virus infections

**DOI:** 10.1101/862037

**Authors:** Zhengchen Su, Qing Chang, Aleksandra Drelich, Thomas Shelite, Barbara Judy, Yakun Liu, Jie Xiao, Changchen Zhou, Xi He, Tais Saito, Shaojun Tang, Lynn Soong, Maki Wakamiya, Xiang Fang, Alexander Bukreyev, Thomas Ksiazek, William K. Russell, Bin Gong

**Author notes:** Correspondence to: Bin Gong, MD, PhD.

## Abstract

Intracerebral microhemorrhages (CMHs) are small foci of hemorrhages in the cerebrum. Acute infections induced by some intracellular pathogens, including rickettsia, can result in CMHs. Annexin a2 (ANXA2) has been documented to play a functional role during intracellular bacterial adhesion. Here we report that *ANXA2*-knockout (KO) mice are more susceptible to CMHs in response to rickettsia and Ebola virus infections, suggesting an essential role of ANXA2 in protecting vascular integrity during these intracellular pathogen infections. Proteomic analysis via mass spectrometry of whole brain lysates and brain-derived endosomes from *ANXA2*-KO and wild-type (WT) mice post-infection with *R. australis* revealed that a variety of significant proteins were differentially expressed, and the follow-up function enrichment analysis had identified several relevant cell-cell junction functions. Immunohistology study confirmed that both infected WT and infected *ANXA2*-KO mice were subjected to adherens junctional protein (VE-cadherin) damages. However, key blood-brain barrier (BBB) components, tight junctional proteins ZO-1 and occludin, were disorganized in the brains from *R. australis*-infected *ANXA2*-KO mice, but not those of infected WT mice. Similar *ANXA2*-KO dependent CMHs and fragments of ZO-1 and occludin were also observed in Ebola virus-infected *ANXA2*-KO mice, but not found in infected WT mice. Overall, our study revealed a novel role of ANXA2 in the formation of CMHs during *R. australis* and Ebola virus infections; and the underlying mechanism is relevant to the role of ANXA2-regulated tight junctions and its role in stabilizing the BBB in these deadly infections.

**Author Summary:** Traditionally, spontaneous intracerebral microhemorrhages (CMHs) were defined as small foci of intracerebral hemorrhages. Such atraumatic CMHs are due to the rupture of a weak blood vessel wall. Infections complicating cerebrovascular accidents have been extensively investigated. However, the role of CMHs complicating infections, in particularly acute systemic infections, has been poorly explored. Population-based retrospective cohort studies suggest there are potentially more undiscovered cases of CMHs accompanying acute systemic infections. Given both the lack of an animal model and cellular/molecular pathophysiology of CMHs following acute systemic infections, there is an urgent need to increase our comprehensive understanding of acute infection-induced CMHs. Overall, our study revealed a novel role of annexin a2 (ANXA2) in the formation of CMHs during *R. australis* and Ebola virus infections; and the underlying mechanism is relevant to the role of ANXA2-regulated endothelial tight junctions and its role in stabilizing the blood-brain barrier in these deadly infections.

## Introduction

Spontaneous intracerebral microhemorrhages (CMHs) are defined as small foci of hemorrhages in the cerebrum [1–3]. These atraumatic CMHs are due to the rupture of small arteries, arterioles, and/or capillaries[4]. CMHs have been a frequently recognized entity since the widespread application of magnetic resonance imaging[1, 5]. Recent investigations into CMHs have seen notable developments, and the increasing prevalence of CMHs is recognized as a significant problem [1, 2]. A population-based retrospective cohort study revealed that 18% of patients with central nervous system (CNS) infections developed CMHs within one year after the initial infection, 47 times greater than non-CNS infection controls[6]. Infections complicating cerebrovascular accidents have been extensively investigated[7–26]. However, the role of CMHs complicating infections[27–30], in particular, acute infections, has been poorly explored[6, 31–33]. To the best of our knowledge, four clinic reports from different countries described a total of 11 cases of CMH after acute systemic infections in patients ranging in age from 9 to 71 years[34–37]. Nine cases in two reports correlated the CMHs with specific pathogen infections, spotted fever (SF) rickettsiosis or ehrlichiosis[37]. Taken together, this information suggests there are potentially more undiscovered cases of CMHs accompanying acute systemic infections.

CMH can be acutely caused by pathogen-associated inflammation (AICMHs)[32, 38]. One of the underlying pathology of AICMHs is the acute dysfunction of the blood-brain barrier (BBB)[39] which is the interface between circulating blood and the central nervous system (CNS) and is composed of brain microvascular endothelial cells (BMECs), pericytes, and astrocytes[40, 41]. BBB properties are primarily determined by junctional complexes between the BMECs, i.e. adherens junctions (AJs) and tight junctions (TJs)[42–44]. Several infectious agents, including rickettsia and Ebola virus, can directly or indirectly target brain endothelial barrier function. Rickettsioses represent devastating human infections[45]. These arthropod-borne diseases are caused by obligate intracellular bacteria of the genus *Rickettsia spp* (*R*.)[46–51]. Disseminated infection of vascular endothelial cell (EC) and endothelial barrier dysfunction are the central pathophysiologic features of human lethal spotted fever group rickettsiosis (SFGR) [45, 46, 52-55]. Ebola virus is associated with severe hemorrhagic diseases in humans. Similar to rickettsia, Ebola virus targets both the immune system[56–63] and ECs, causing severe vascular leakage syndrome[64–68], but the underlying mechanisms remain unclear. Two main mechanisms have been proposed: the virus infects macrophages and dendritic cells, which then release cytokines to activate ECs; and/or the virus directly infects and activates ECs[67, 69].

Annexin A2 (ANXA2) is a member of the large annexin family of Ca^2+^-regulated and phospholipid-binding proteins, which associates with cell membrane dynamics, cell-cell interactions, and cell adhesion[70–76]. ANXA2 can be monomeric, found mainly in the cytosol, or forming heterotetramer complex with S100A10 [72, 77]. S100A10 is a unique member of S100 protein family that has been known to bind to ANXA2. The interaction between ANXA2 and S100A10 yields a heterotetramer complex ANXA2-S100A10, enabling ANXA2 to translocate across the EC membrane and perform a variety of functions, facilitating plasmin-based fibrinolytic activities on vascular luminal surfaces[71, 72, 78]. Recently, we identified host ANXA2 as a novel receptor for SFGR and staphylococcus aureus adhesions to ECs[79]. However, there was no difference in rickettsial adhesion to or invasion into white blood cells between the wild-type (WT) and *ANXA2*-knockout (KO) mice.

Here we report an observation that focal CMH lesions exist in the cerebra of *R. australis* infected *AXNA2*-KO mice but not *R. australis* infected WT mice. We hypothesize cell-cell junction in the BBB is destabilized in *AXNA2*-KO mice rendering them susceptible to AICMH. In order to comprehensively investigate this possibility, we performed a proteomic analysis using the whole brain lysate and brain-derived isolated endosomes. We identified a variety of differentially expressed (DE) proteins that were relevant to vascular integrity. Functional group annotation and network analysis based on the identified DE proteins revealed a variety of protein functional group changes, such as cell-cell junction, stress fiber, MHC II protein complex binding, and stress response. These identified functional groups support that a structural impairment of the BBB might be involved. Consistently, immunofluorescence (IF) of brain tissue of *R. australis* infected mice revealed dramatic disruption and disorganization of TJ proteins ZO-1 and occludin in *ANXA2*-KO mice, but not WT mice. Interestingly, *ANXA2*-KO mice challenged by Ebola virus also exhibited CMHs and aberrant TJs whereas WT mice showed no signs of bleeding into CNS, indicating this pathology is not specific to rickettsia infection. Collectively, these data suggest that ANXA2 is required for the integrity of TJs in response to acute rickettsia and Ebola infections.

## Results

### 1. Absence of ANXA2 is associated with the incidence of CMH in *R. australis* infection

To investigate the potential role of *ANXA2*-KO on survival of the mice in response to the *R. australis* infection, WT (n=14) and *ANXA2*-KO (n=15) mice were inoculated with an ordinary lethal dose of *R. australis* (2 x 10^6^) via tail vein injection (i.v.) [79, 80] and observed up to 10 days post-infection (p.i.). Accumulative survival data were obtained from three independent experiments (**Supplemental Fig. 1A**) and subjected to Kaplan-Meier (K-M) analysis. We found no difference in survival between WT (21.43%) and *ANXA2*-KO (13.33%) groups. However, the gross pathology of the brain surface (**Supplemental Fig. 1B**) observed apparent different color cerebral areas between infected WT mice and infected *ANXA2*-KO. To examine underlying associated pathology(s), brain tissue sections were subjected to histological examination with hematoxylin and eosin (H&E) staining (**Fig. 1**), which revealed striking focal CMHs, in the cerebra of all lethally infected *ANXA2*-KO mice (**Fig. 1G-L**), but not in the two survival infected *ANXA2*-KO mice. Conversely, such CMHs were absent from both lethal and survival WT infected mice (**Fig. 1 D-F**).

**Figure 1.**
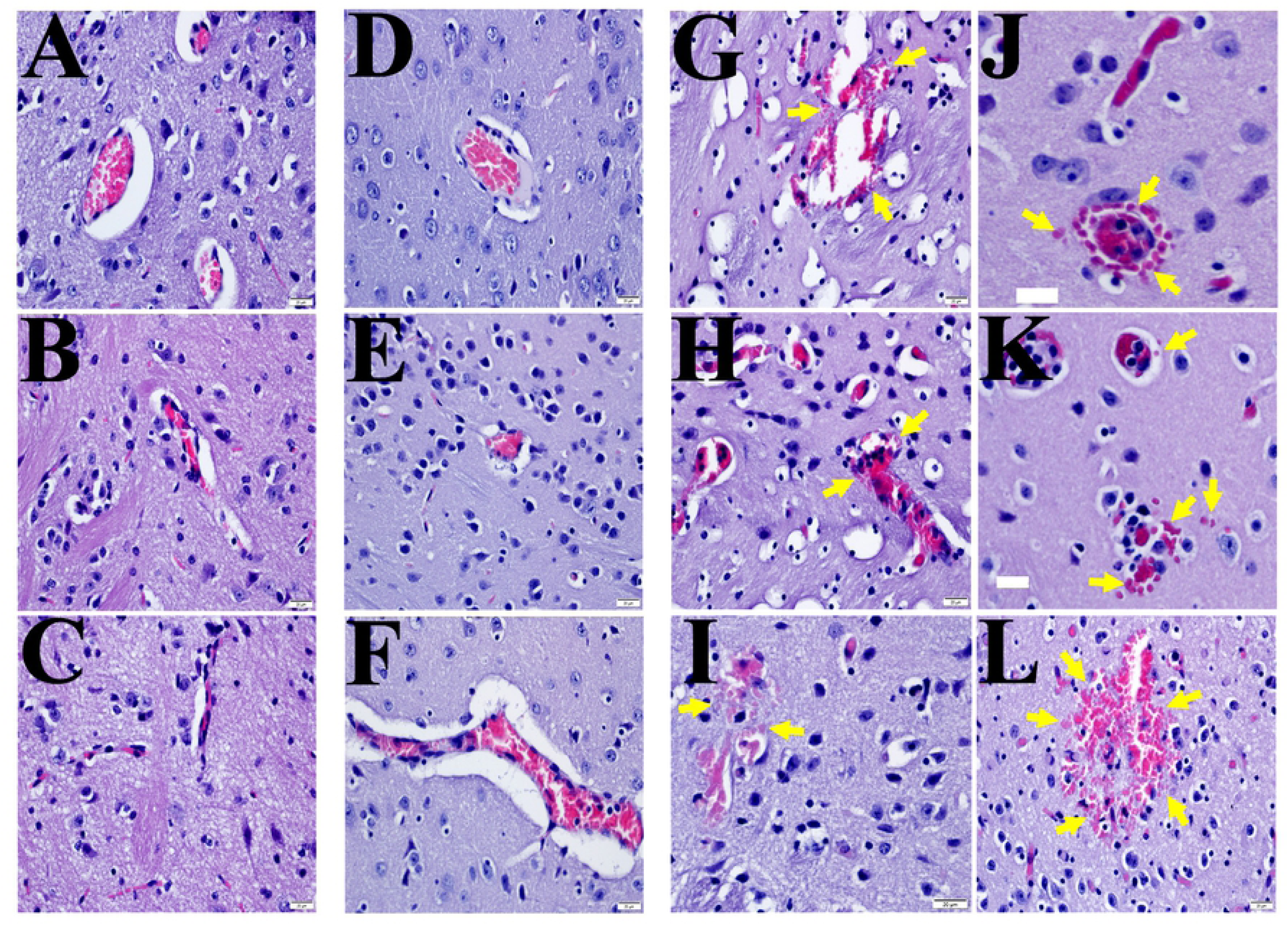
Representative H&E staining of the brains from *R.australlis*-infected *ANXA2*-KO and WT mice. Yellow arrows indicate the presence of focal hemorrhagic lesions. A, mock WT; B&C, mock *AXNA2*-KO; D-F, *R. australis*-infected WT; G-L), *R. australis*-infected *AXNA2*-KO mice. Scale bar: 20 µm.

### 2. Inactivation of ANXA2 does not affect the proliferation of *R. australis* and serum levels of IFNγ and TNFα in mice

For time-dependent pathological studies, mice were inoculated with an ordinary lethal dose of *R. australis* (2 x 10^6^). At 5 days p.i. (5 mice per group), both *ANXA2*-KO and WT mice were euthanized as designed. H&E examination revealed extensive focal hemorrhagic lesions, at levels of arteriole, capillary, and venule, in the cerebra of all infected *ANXA2*-KO mice on day 5 p.i., but not WT mice (**Fig. 2A-D**).

**Figure 2.**
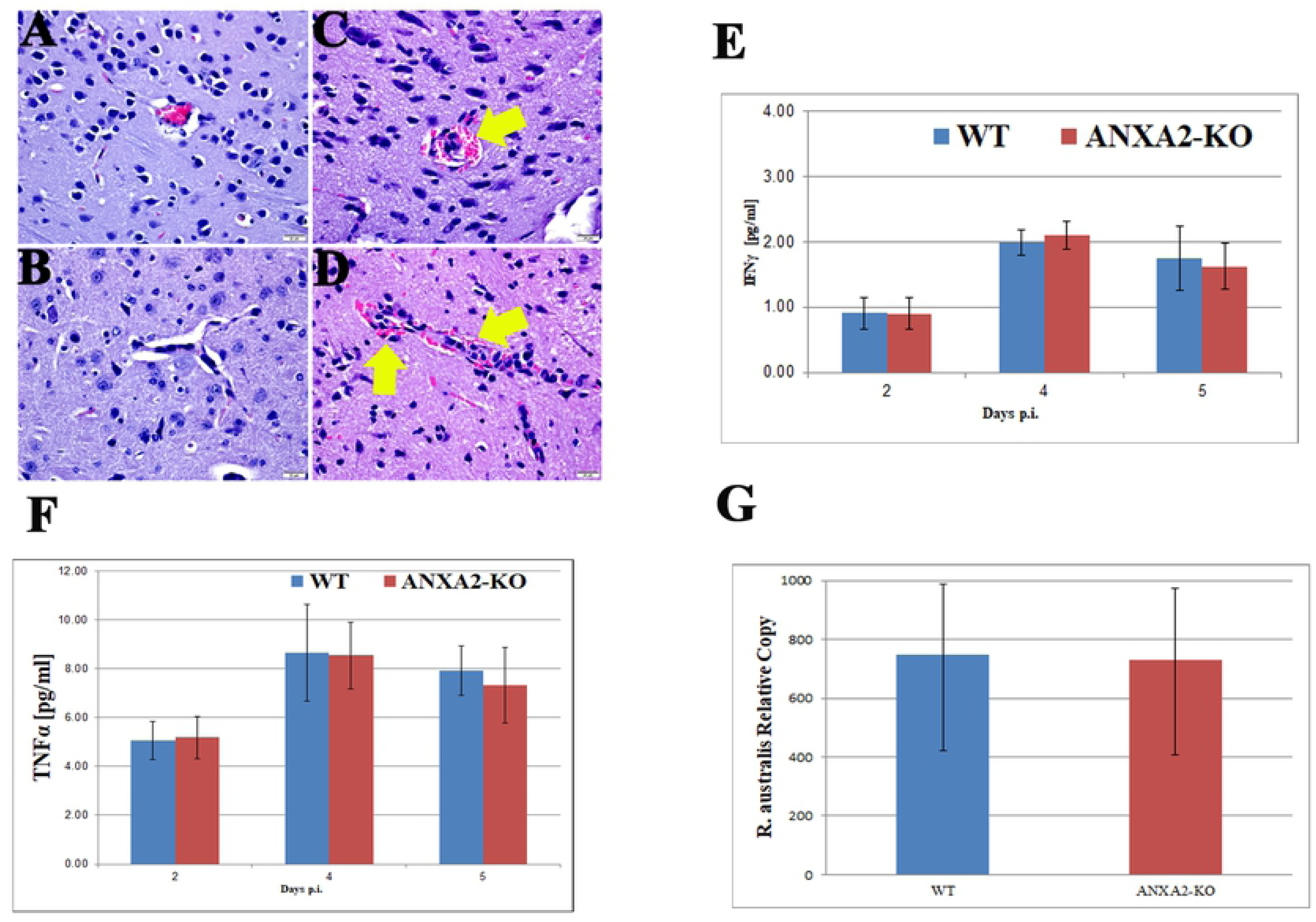
Representative H&E staining of brain sections from WT (A&B) and *ANXA2*-KO (C&D) 5 days post-*R. australis* infection. Perivascular hemorrhage (yellow arrow) can be observed in infected *ANXA2*-KO group but not infected WT group. TNFα (E) and IFNγ (F) concentrations in serum at 2,4,5 days post-*R. australis* infection. Relative R. australis DNA copies (G) extracted from the brain of WT and *ANXA2*-KO mice quantified by rt-qPCR. No significant difference was found. Error bar stands for standard deviation. Scale bar: 20 µm.

To examine whether ANXA2 plays a role in serum levels of IFNγ and TNFα during the lethal dose of *R. australis* infection, serum from mice at day 2, 4, or 5 p.i. were collected and proceeded to analyze the concentration of IFNγ and TNFα. However, no difference was found comparing WT and *ANXA2*-KO mice (**Fig. 2E and F**). Furthermore, on day 5 post-infection, the real-time qPCR analysis revealed no difference in bacterial loads in brain between WT (n=4) and *AXNA2*-KO mice (n=4) (**Fig. 2G**). Immunofluorescent staining (IF) of rickettsia in the liver, brain and lung did not show any difference between WT and *ANXA2*-KO mice on day 5 p.i. (**Fig. 3**). These data suggest *ANXA2* depletion does not affect the overall proliferation of *R. australis* in the mice.

**Figure 3.**
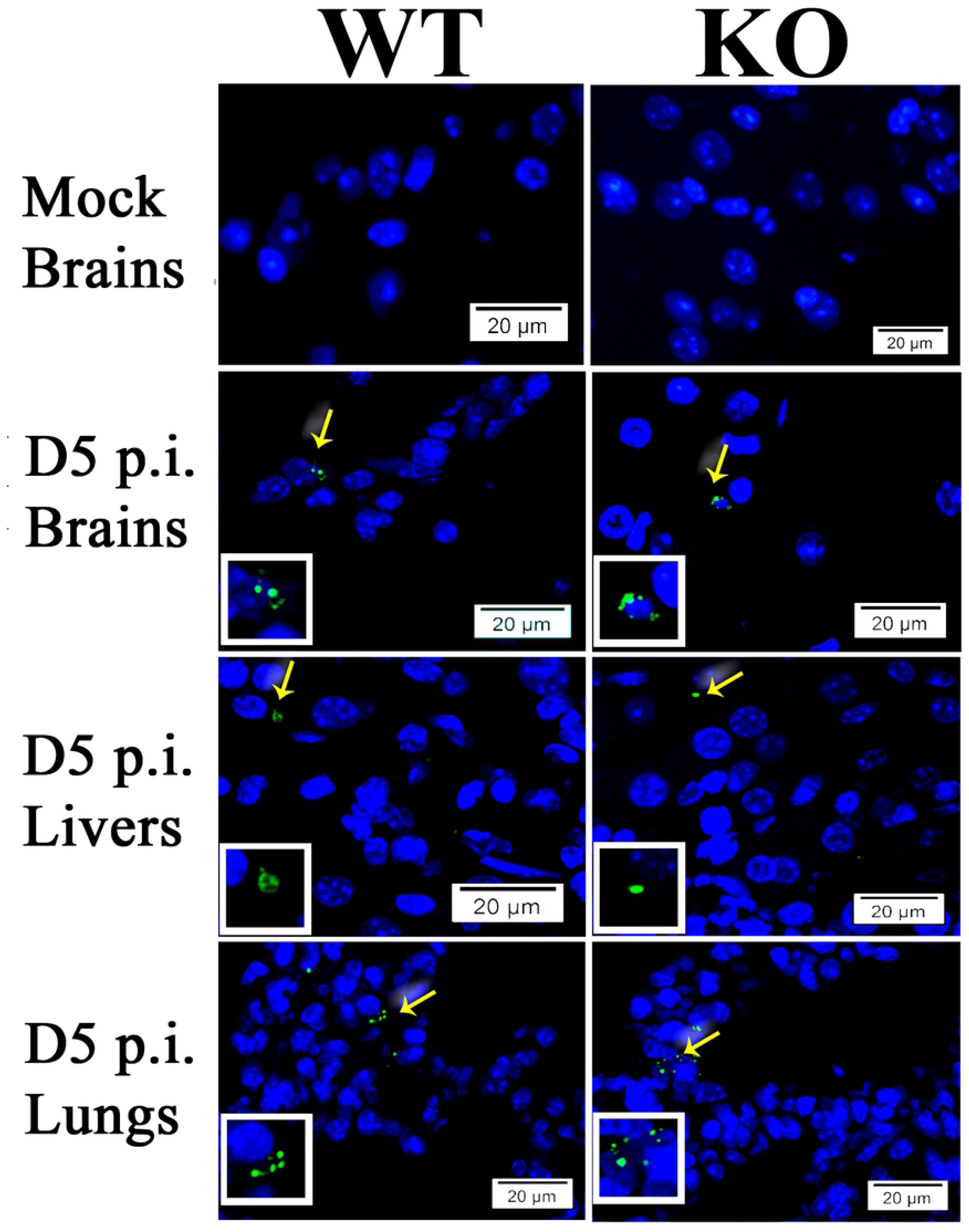
Representative IF staining of SFG rickettsiae (green) in livers, brains and lungs from WT and *ANXA2*-KO mice while nuclei of mouse cells were counter-stained with DAPI (blue). The areas indicated by the arrows are enlarged and distinguish rickettsial (green) staining (boxed inserts). Scale bars, 20 µm.

### 3. Proteomic analysis

In order to decipher the protein profile related to the CMHs observed in the *ANXA2-*KO mice post rickettsia infection, we performed a proteomic analysis of the whole brain protein lysate and isolated endosome. Isolated endosome protein pattern is important due to the involvement of ANXA2 in endocytosis and turnover of surface proteins and nucleotide[72].

#### 3.1 Whole-brain lysate

Comparing the protein profiles of brain lysate from *ANXA2*-KO and WT mice challenged by *R.australis* infection on day 5 p.i., LC/MS analysis identified one hundred thirty DE, with 93 upregulated and 37 downregulated (*ANXA2*-KO versus WT). The top upregulated and downregulated proteins are listed in **Tables 1 & 2**. It is noteworthy that the hemoglobin level was higher (4-fold) in the infected *ANXA2-*KO mice, suggesting to be the cause of different gross pathology from the infected WT mice. Functional enrichment analysis and visualization (**Fig. 4 & 6**) revealed statistically significant functional groups that are potentially associated with CMH in *ANXA2-*KO mice. The noteworthy upregulated proteins that are associated with the cell junction structure integrity include heat shock protein 90 α (HSP90α), heat shock protein 90 beta, enolase 1, clathrin (heavy polypeptide); important downregulated proteins include myosin (heavy polypeptide 10).

**Figure 4.**
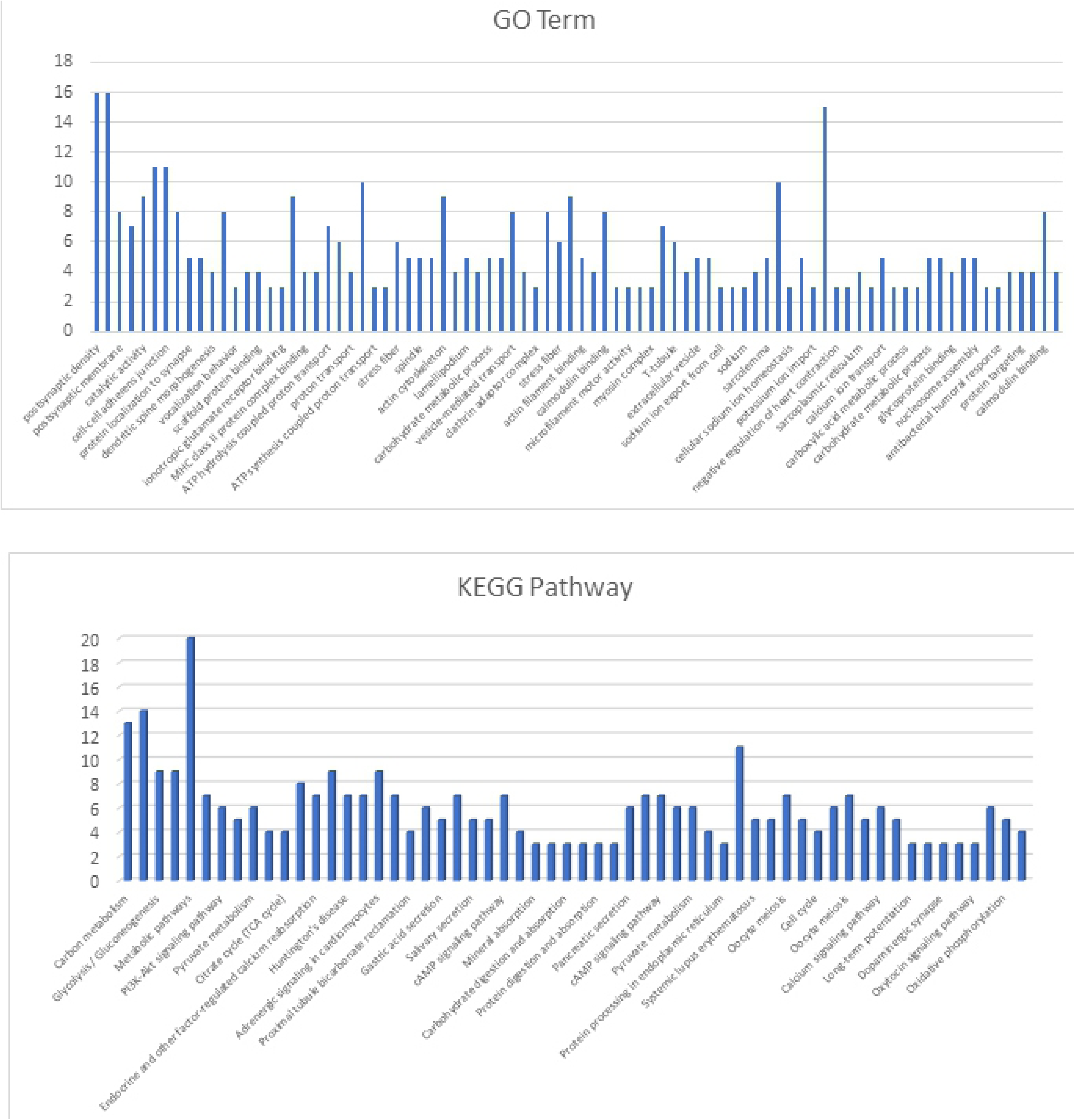
Functional annotation from Go Term and KEGG for whole-brain lysate.

**Figure 5.**
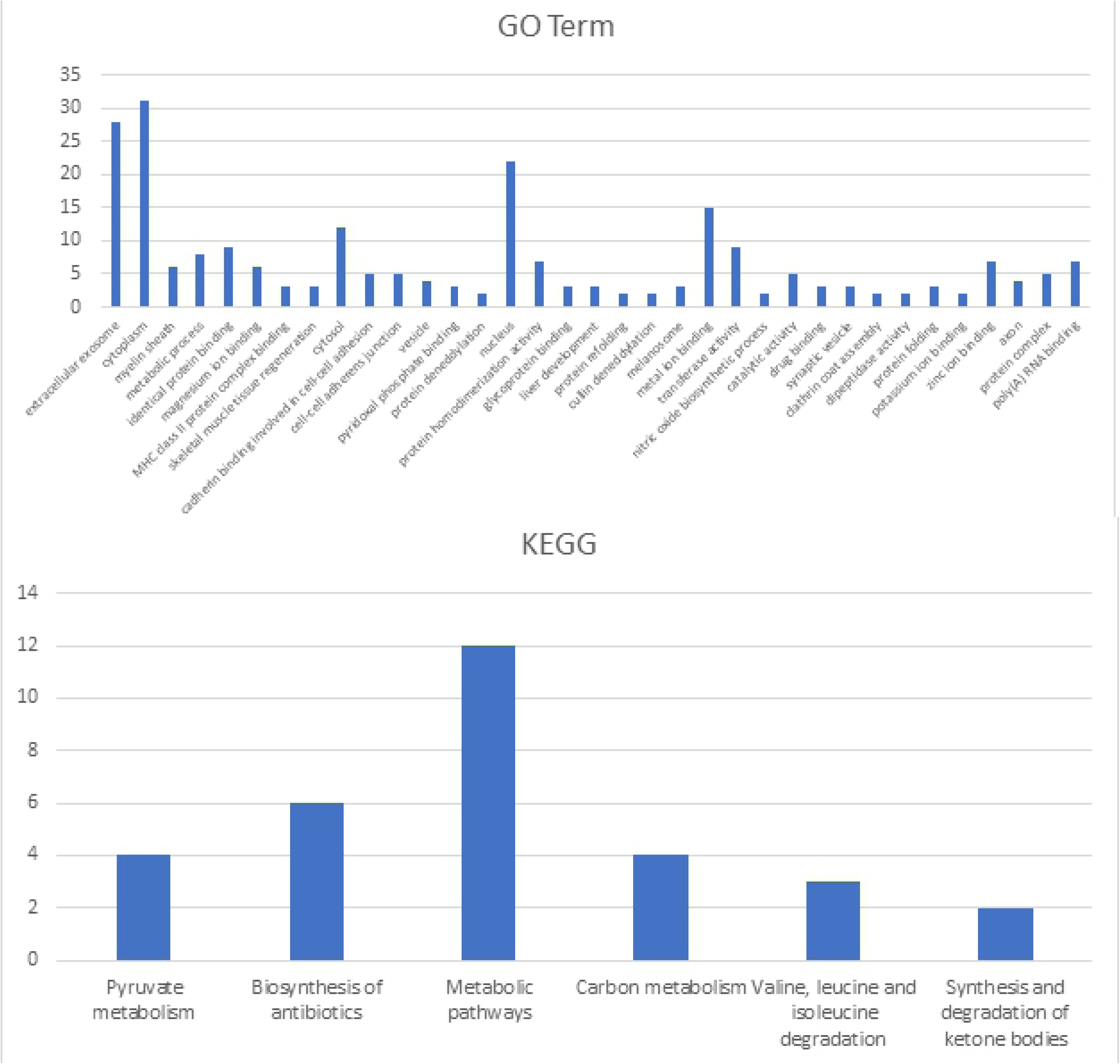
Functional annotation from Go Term and KEGG for isolated endosomes from the brain.

**Figure 6.**
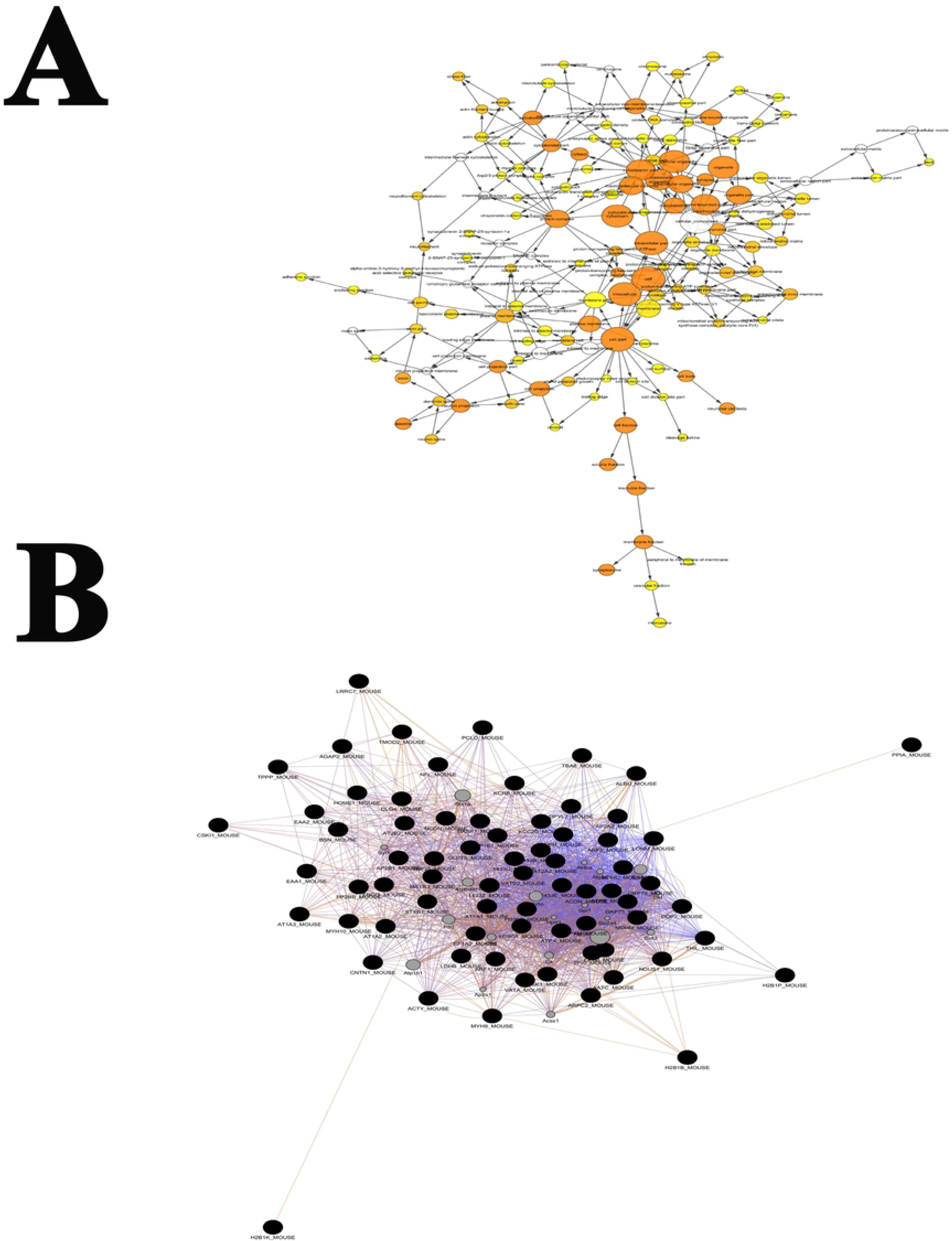
Functional group from GO Term Cellular Component (A) and DE protein-protein interaction network visualization (B) for whole-brain lysate.

**Table 1.** Upregulated proteins and their KEGG pathway interpretation from whole-brain lysate. Fold change is calculated as (KO-WT)/WT.

| Gene ID | Protein | Fold change | KEGG Pathway |
| --- | --- | --- | --- |
| AT2B1_MOUSE | ATPase, Ca<sup>++</sup> transporting, plasma membrane 1(Atp2b1) | 4.17 | Calcium signaling pathway, cGMP-PKG signaling pathway, cAMP signaling pathway, Adrenergic signaling in cardiomyocytes, Salivary secretion, Pancreatic secretion, |
| Q3UHH0_MOUSE, AT2B2_MOUSE | ATPase, Ca<sup>++</sup> transporting, plasma membrane 2(Atp2b2) | 6.25 | Calcium signaling pathway, cGMP-PKG signaling pathway, cAMP signaling pathway, Adrenergic signaling in cardiomyocytes, Salivary secretion, Pancreatic secretion, |
| VATA_MOUSE | ATPase, H<sup>+</sup> transporting, lysosomal V1 subunit A(Atp6v1a) | 4.36 | Oxidative phosphorylation, Metabolic pathways, Phagosome, Synaptic vesicle cycle, Collecting duct acid secretion, Rheumatoid arthritis, |
| AP1B1_MOUSE | adaptor protein complex AP-1, beta 1 subunit(Ap1b1) | 5.00 | Lysosome, |
| CLH1_MOUSE | clathrin, heavy polypeptide (Hc)(Cltc) | 5.83 | Lysosome, Endocytosis, Synaptic vesicle cycle, Endocrine and other factor-regulated calcium reabsorption, Huntington's disease, Bacterial invasion of epithelial cells, |
| ENOA_MOUSE | enolase 1, alpha non-neuron(Eno1) | 4.33 | Glycolysis / Gluconeogenesis, Metabolic pathways, Biosynthesis of antibiotics, Carbon metabolism, Biosynthesis of amino acids, RNA degradation, HIF-1 signaling pathway, |
| HS90B_MOUSE | heat shock protein 90 alpha (cytosolic), class B member 1(Hsp90ab1) | 4.10 | Protein processing in endoplasmic reticulum, PI3K-Akt signaling pathway, Antigen processing and presentation, NOD-like receptor signaling pathway, Progesterone-mediated oocyte maturation, Estrogen signaling pathway, Pathways in cancer, Prostate cancer, |
| **HS90A_MOUSE** | heat shock protein 90, alpha (cytosolic), class A member 1(Hsp90aa1) | 9.40 | Protein processing in endoplasmic reticulum, PI3K-Akt signaling pathway, Antigen processing and presentation, NOD-like receptor signaling pathway, Progesterone-mediated oocyte maturation, Estrogen signaling pathway, Pathways in cancer, Prostate cancer, |
| **A8DUK4_MOUSE** | hemoglobin, beta adult s chain(Hbb-bs) | 4.09 | African trypanosomiasis, Malaria, |
| **H2B1B_MOUSE** | histone cluster 1, H2bb(Hist1h2bb) | 4.67 | Alcoholism, Viral carcinogenesis, Systemic lupus erythematosus, |
| **H2B1K_MOUSE** | histone cluster 1, H2bk(Hist1h2bk) | 4.67 | Alcoholism, Viral carcinogenesis, Systemic lupus erythematosus, |
| **Q8CBB6_MOUSE** | histone cluster 1, H2bq(Hist1h2bq) | 4.67 | Alcoholism, Viral carcinogenesis, Systemic lupus erythematosus, |
| **MDHM_MOUSE** | malate dehydrogenase 2, NAD (mitochondrial)(Mdh2) | 11.75 | Citrate cycle (TCA cycle), Cysteine and methionine metabolism, Pyruvate metabolism, Glyoxylate and dicarboxylate metabolism, Metabolic pathways, Biosynthesis of antibiotics, Carbon metabolism, |
| **PFKAM_MOUSE** | phosphofructokinase, muscle(Pfkm) | 4.25 | Glycolysis / Gluconeogenesis, Pentose phosphate pathway, Fructose and mannose metabolism, Galactose metabolism, Metabolic pathways, Biosynthesis of antibiotics, |
| **KPYM_MOUSE** | pyruvate kinase, muscle(Pkm) | 8.83 | Glycolysis / Gluconeogenesis, Purine metabolism, Pyruvate metabolism, Metabolic pathways, Biosynthesis of antibiotics, Carbon metabolism, Biosynthesis of amino acids, |
| **SNP25_MOUSE** | synaptosomal-associated protein 25(Snap25) | 4.33 | Synaptic vesicle cycle, Insulin secretion, |
| **1433E_MOUSE** | tyrosine 3-monooxygenase/tryptophan 5-monooxygenase activation protein, epsilon polypeptide(Ywhae) | 4.17 | Cell cycle, Oocyte meiosis, PI3K-Akt signaling pathway, Hippo signaling pathway, Neurotrophin signaling pathway, Viral carcinogenesis, |

**Table 2.** Downregulated proteins and their KEGG pathway interpretation from whole-brain lysate. Fold change is calculated as (KO-WT)/WT.

| **Gene ID** | **Protein name** | **Fold change** | **Representative KEGG Pathway** |
| --- | --- | --- | --- |
| **A4GZ26_MOUSE, E9QAD8_MOUSE, D3Z5I6_MOUSE** | IQ motif and Sec7 domain 2(Iqsec2) | -0.63 | Endocytosis, |
| **D3YZU5_MOUSE** | SH3/ankyrin domain gene 1(Shank1) | -0.7 | Glutamatergic synapse, |
| **KCC2G_MOUSE** | calcium/calmodulin-dependent protein kinase II gamma(Camk2g) | -0.63 | ErbB signaling pathway, Calcium signaling pathway, cAMP signaling pathway, HIF-1 signaling pathway, |
| **E9Q1T1_MOUSE, A0A0G2JGS4_MOUSE** | calcium/calmodulin-dependent protein kinase II, delta(Camk2d) | -0.76 | ErbB signaling pathway, Calcium signaling pathway, cAMP signaling pathway, HIF-1 signaling pathway, |
| **MYH10_MOUSE, Q5SV64_MOUSE, Q3UH59_MOUSE** | myosin, heavy polypeptide 10, non-muscle(Myh10) | -0.78 | Tight junction, |
| **PCLO_MOUSE** | piccolo (presynaptic cytomatrix protein)(Pclo) | -0.76 | Insulin secretion, |

Functional enrichment analysis was performed based on Gene Ontology (GO) Term and Kyoto Encyclopedia of Genes and Genomes (KEGG) pathway. Noteworthy GO Term include *cell-cell adherens junction, stress fiber, actin cytoskeleton, MHC class II protein complex binding, vesicle-mediated transport, and extracellular vesicle.* In KEGG pathway, we identified a variety of signaling pathways, in which *PI3K-Akt signaling pathway, cAMP signaling pathway, and protein digestion and absorption pathway* may be relevant to the pathology of CMHs. The potential interacting network of DE proteins is shown in **Fig. 6**.

#### 3.2 Brain-derived isolated endosome

The endosome is integral to the endocytosis pathway. Due to the nature of endosomes, the proteins enriched in the endosomes is either subjected to degradation or recycled back to the plasma membrane[81]. Increased accumulation of proteins in the endosomal compartment suggests increased protein turnover rate and degradation; whereas reduction of protein enrichment in the endosomes might result in protein accumulation in another compartment such as cytosol or plasma membrane[81].

ANXA2 is known to be associated with the endosomal membrane. Endocytosis of several targets depends on the presence of tyrosine phosphorylation of ANXA2[82–87]. The absence of ANXA2 may lead to a differential endosomal protein profile, which may shed light on the underlying mechanisms associated with the increased susceptibility of *ANXA2*-KO mice to CMH upon rickettsia infection.

Comparing the endosomes isolated from the mouse brains of WT mice and *ANXA2*-KO 5 days after rickettsial infection, we have identified 47 DE proteins, with 18 upregulated proteins, and 29 downregulated proteins in the infected *ANXA2*-KO mice. Noteworthy upregulated DE proteins include talin1, alanyl-tRNA synthetase, glutathione peroxidase 1, and calcium/calmodulin-dependent protein kinase IV. Important downregulated proteins are cofilin 2, heat shock protein 8, HSP90α, thimet oligopeptidase and seryl-aminoacyl-tRNA synthetase. Top proteins are listed in **Table 3 & 4**. Functional enrichment analysis was performed based on these 47 DE proteins, generating a list of functional enriched clusters (**Fig. 5 & 7**). Noteworthy Go Term functional groups include cell-cell AJ, metalloproteinase, and MHC II protein complex binding. Significantly altered signaling pathways in KEGG are pyruvate metabolism, biosynthesis of antibiotics, metabolic pathways, caron metabolism, valine&leucine&isoleucine degradation, synthesis and degradation of ketone bodies.

**Figure 7.**
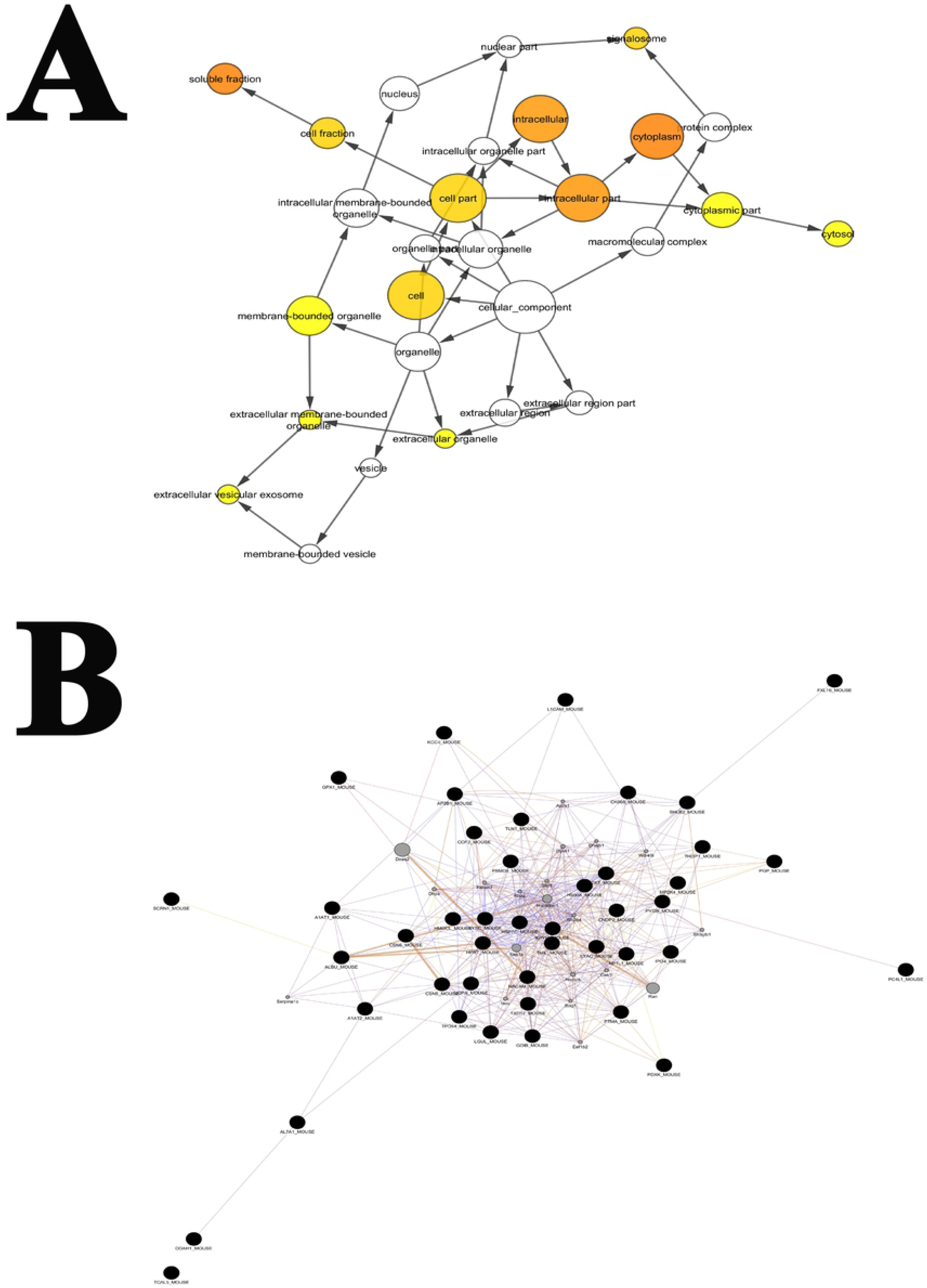
Functional group from GO Term Cellular Component (A) and DE protein-protein interaction network visualization (B) for brain-derived endosome.

**Table 3.** Upregulated proteins and their KEGG pathway interpretation from brain-derived endosomes. Fold change is calculated as (KO-WT)/WT.

| ID | Gene Name | fold change | KEGG_PATHWAY |
| --- | --- | --- | --- |
| **HMGCL_MOUSE** | 3-hydroxy-3-methylglutaryl-Coenzyme A lyase(Hmgcl) | 2 | Synthesis and degradation of ketone bodies, Valine, leucine and isoleucine degradation, Butanoate metabolism, Metabolic pathways, Peroxisome, |
| **L1CAM_MOUSE** | L1 cell adhesion molecule(L1cam) | 2.5 | Axon guidance, Cell adhesion molecules (CAMs), |
| **SHLB2_MOUSE** | SH3-domain GRB2-like endophilin B2(Sh3glb2) | 2.33 | Endocytosis, |
| **AP2B1_MOUSE** | adaptor-related protein complex 2, beta 1 subunit(Ap2b1) | 3.5 | Endocytosis, Synaptic vesicle cycle, Endocrine and other factor-regulated calcium reabsorption, Huntington's disease, |
| **SYAC_MOUSE** | alanyl-tRNA synthetase(Aars) | 2.17 | Aminoacyl-tRNA biosynthesis, |
| **PYGB_MOUSE** | brain glycogen phosphorylase(Pygb) | 4.5 | Starch and sucrose metabolism ,Metabolic pathways, Insulin signaling pathway, Glucagon signaling pathway,Insulin resistance, |
| **KCC4_MOUSE** | calcium/calmodulin -dependent protein kinase IV(Camk4) | 2 | Calcium signaling pathway, cAMP signaling pathway, Osteoclast differentiation, Long-term potentiation, Neurotrophin signaling pathway, Cholinergic synapse, Oxytocin signaling pathway, Aldosterone synthesis and secretion, Amphetamine addiction, Alcoholism, |
| **GPX1_MOUSE** | glutathione peroxidase 1(Gpx1) | 3 | Glutathione metabolism, Arachidonic acid metabolism,Thyroid hormone synthesis, |
| **PGP_MOUSE** | phosphoglycolate phosphatase(Pgp) | 3 | Glyoxylate and dicarboxylate metabolism, Metabolic pathways, Biosynthesis of antibiotics, Carbon metabolism, |
| **TLN1_MOUSE** | talin 1(Tln1) | 3.33 | Rap1 signaling pathway, Focal adhesion, Platelet activation, HTLV-I infection, |

**Table 4.** Downregulated proteins and their KEGG pathway interpretation from brain-derived endosomes. Fold change is calculated as (KO-WT)/WT.

| ID | Gene Name | Fold change | Representative KEGG_PATHWAY |
| --- | --- | --- | --- |
| **CNDP2_MOUSE** | CNDP dipeptidase 2 (metallopeptidase M20 family)(Cndp2) | -0.6 | Arginine and proline metabolism, Histidine metabolism, beta-Alanine metabolism, Metabolic pathways, |
| **UGPA_MOUSE** | UDP-glucose pyrophosphorylase 2(Ugp2) | -0.75 | Pentose and glucuronate interconversions, Galactose metabolism, Starch and sucrose metabolism, Amino sugar and nucleotide sugar metabolism, Metabolic pathways, Biosynthesis of antibiotics, |
| **THIL_MOUSE** | acetyl-Coenzyme A acetyltransferase 1(Acat1) | -0.67 | Fatty acid degradation, Synthesis and degradation of ketone bodies, Valine, leucine and isoleucine degradation, Lysine degradation, Tryptophan metabolism, Pyruvate metabolism, |
| **AL7A1_MOUSE** | aldehyde dehydrogenase family 7, member A1(Aldh7a1) | -0.71 | Glycolysis / Gluconeogenesis, Ascorbate and aldarate metabolism, Fatty acid degradation, Glycine, serine and threonine metabolism, Valine, leucine and isoleucine degradation, Lysine degradation, |
| **BHMT1_MOUSE** | betaine-homocysteine methyltransferase(Bhmt) | -0.67 | Glycine, serine and threonine metabolism, Cysteine and methionine metabolism, Metabolic pathways, |
| **COF2_MOUSE** | cofilin 2, muscle(Cfl2) | -0.63 | Axon guidance, Fc gamma R-mediated phagocytosis, Regulation of actin cytoskeleton, Pertussis, |
| **LGUL_MOUSE** | glyoxalase 1(Glo1) | -0.62 | Pyruvate metabolism, |
| **HSP7C_MOUSE** | heat shock protein 8(Hspa8) | -0.73 | Spliceosome, MAPK signaling pathway, Protein processing in endoplasmic reticulum, Endocytosis, Antigen processing and presentation, Estrogen signaling pathway, Legionellosis, Toxoplasmosis, Measles, Influenza A, |
| **HS90A_MOUSE** | heat shock protein 90, alpha (cytosolic), class A member 1(Hsp90aa1) | -0.63 | Protein processing in endoplasmic reticulum, PI3K-Akt signaling pathway, Antigen processing and presentation, NOD-like receptor signaling pathway |
| **HPRT_MOUSE** | hypoxanthine guanine phosphoribosyl transferase(Hprt) | -0.67 | Purine metabolism, Drug metabolism - other enzymes, Metabolic pathways, |
| **PDXK_MOUSE** | pyridoxal (pyridoxine, vitamin B6) kinase(Pdxk) | -0.78 | Vitamin B6 metabolism, Metabolic pathways, |
| **KPYM_MOUSE** | pyruvate kinase, muscle(Pkm) | -0.67 | Glycolysis / Gluconeogenesis, Purine metabolism, Pyruvate metabolism, Metabolic pathways, Biosynthesis of antibiotics, Carbon metabolism, Biosynthesis of amino acids, |
| **A1AT1_MOUSE** | serine (or cysteine) peptidase inhibitor, clade A, member 1A(Serpina1a) | -0.64 | Complement and coagulation cascades, |
| **A1AT2_MOUSE** | serine (or cysteine) peptidase inhibitor, clade A, member 1B(Serpina1b) | -0.64 | Complement and coagulation cascades, |
| **SYSC_MOUSE** | seryl-aminoacyl-tRNA synthetase(Sars) | -0.69 | Aminoacyl-tRNA biosynthesis, |
| **THOP1_MOUSE** | thimet oligopeptidase 1(Thop1) | -0.78 | Renin-angiotensin system, African trypanosomiasis, |
| **TKT_MOUSE** | transketolase(Tkt) | -0.83 | Pentose phosphate pathway, Metabolic pathways, Biosynthesis of antibiotics, Carbon metabolism, Biosynthesis of amino acids, |

### 4. Endothelial TJ proteins disorganized in brains of *ANXA2*-KO mice after lethal SFGR infections

The overall functional enrichment analysis supports our hypothesis that deletion of ANXA2 affects the cell-cell junction structure during the rickettsial infection. However, LC/MS showed that the expression level of TJ protein ZO-1 in the brain was similar between infected *ANXA2*-KO and infected WT mice. We postulated that strutures of TJ might be disorganized instead of downregulation of the expression levels.

Occludin, directly interacted with ZO-1, is an important transmembrane protein integral to TJs[88]. To investigate the structure of TJs, we applied IF assay to the brain tissue to examine ZO-1 and occludin. Remarkably, in all five *ANXA2-*KO mice, dramatic disruption and disorganization of ZO-1 and occludin were detected after D5 p.i. (**Fig. 8A**). Interestingly, occludin disorganization was also observed in the livers of these *ANXA2*-KO mice on day 5 p.i. (**Fig. 8B**). AJ protein VE-cadherins were also investigated. Animals in both WT and *ANXA2*-KO groups exhibited similar disintegrated structures of VE-cadherin in response to *R. australis* infection on day 5 p.i. (**Fig. 8C**).

**Figure 8.**
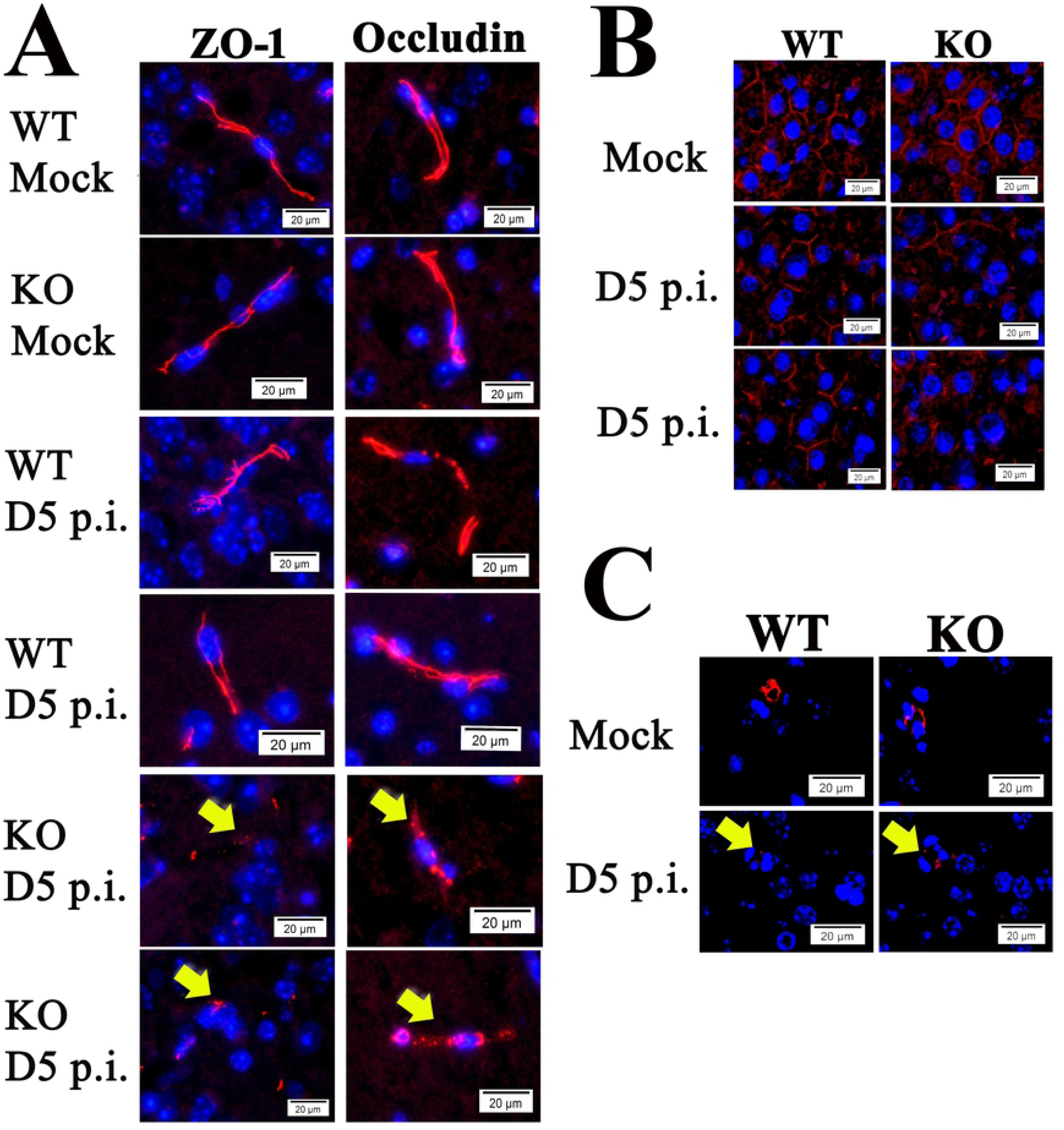
(A) Representative IF staining of ZO-1 (left) and occludin (right) in the brains from *R.australis*/mock-infected/ WT and *ANXA2*-KO mice, at day 5 p.i.. Yellow arrows indicate the fragmented structures of ZO-1 and occludin, which were mainly seen in infected *ANXA2*-KO mice. (B) Representative IF staining for occludin in the livers from *R.australis*/mock-infected/ WT and *ANXA2*-KO mice, at day 5 p.i.. (C) Representative IF for VE-cadherin in the brains from *R.australis*/mock-infected/ WT and *ANXA2*-KO mice, at day 5 p.i. Yellow arrows indicate the fragmented structures of VE-cadherin. Nuclei of mouse cells were counterstained with DAPI (blue). Scale bar: 20um.

### 5. CMHs and disorganized EC TJs in ANXA2-KO mice subjected to Ebola virus infection

Similar to rickettsia, Ebola virus is known to attack the endothelial cells and a severe vascular leakage syndrome is a major complication during Ebola infection[64–69]. We investigated whether the *ANXA2* deletion-dependent CMHs was also present in Ebola virus infection via retrospective histology examination of brain tissue samples from WT (n=5) or *ANXA2*-KO (n=5) mice challenged by a mouse-adapted strain of Ebola Zaire virus (50 plaque-forming unit by the intraperitoneal route[89]). A 20% cumulative survival was observed in *ANXA2*-KO mice 12 days p.i., compared with an 80% cumulative survival in the group of WT mice at the same time (p=0.08, Log-rank test) (**Supplemental Fig. 2**). IF assay of Ebola virus in the liver, brain and lung tissues showed no difference between WT and *ANXA2*-KO mice on day 7-12 p.i. (**Fig. 9A**).

**Figure 9.**
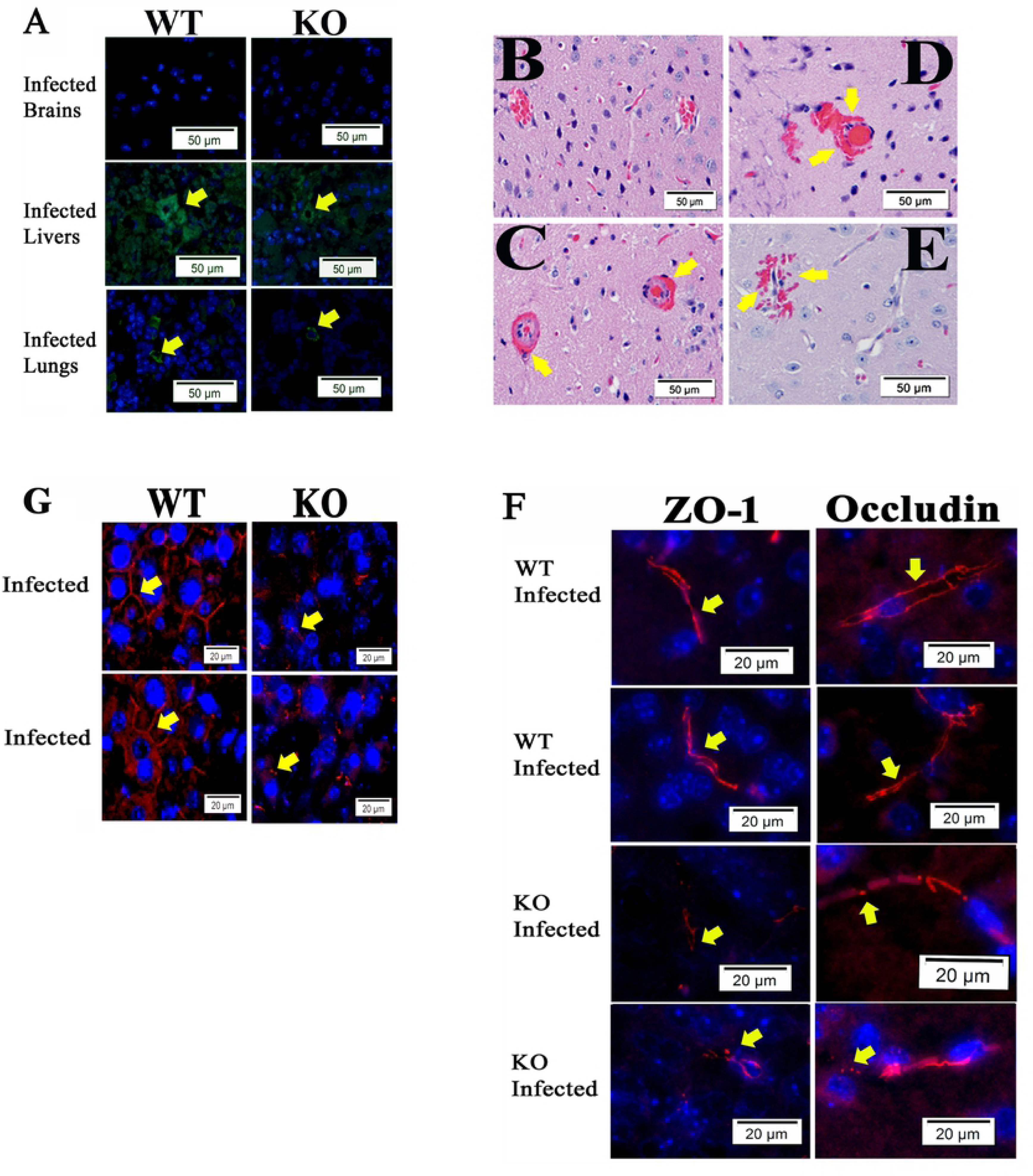
(A) Representative IF for Ebola virus antigen (Green) in the brains, livers, and lungs from WT or *ANXA2*-KO mice post-Ebola virus challenge. Yellow arrows represent the presence of Ebola virus antigen. (B-E) H&E staining of the brain from WT(B) and *ANXA2*-KO (C-E) mice infected by Ebola virus. Yellow arrows indicate the presence of perivascular hemorrhage. (F) Representative IF for ZO-1 (left) and Occludin (right) in the brains from WT or *ANXA2*-KO mice post-Ebola virus challenge. Yellow arrows indicate the position of ZO-1 or occludin, which represent the tight junction structure. *ANXA2*-KO but not WT Ebola virus-infected mice exhibited the fragmented tight junction structure in the brain. (G) Representative IF for occludin (red) in the livers from WT or *ANXA*2-KO mice day 10 post Ebola virus challenge. Yellow arrows show the structure of the pericellular occludin, which was relatively intact in Ebola virus-infected WT mice but dramatically fragmented in the Ebola virus-infected *ANXA2-KO* mice. Nuclei of mouse cells were counterstained with DAPI (blue). Scale bar: 50 µm (A-E), 20 µm (F and G).

Similarly, CMHs was detected in H&E staining in all *ANXA2*-KO mice (**Fig. 9C-E**), but not in WT mice (**Fig. 9B**) during 7-12 days p.i.. Furthermore, disorganized TJ protein ZO-1 and occludin were also present in brain tissue from *ANXA2*-KO, but not WT mice (**Fig. 9F**) infected with same dose of Ebola virus intraperitoneally. Furthermore, dramatic disruption and disorganization of occludin were also observed in the livers of *ANXA2*-KO mice post Ebola virus infection (**Fig. 9G**).

Collectively, these data suggest the detriments to the TJ structures as an underlying mechanism of the CMHs observed in *ANXA2*-KO mice post-*R. australis* or Ebola virus infections.

### 6. Other considerations

An earlier study reported that the deletion of ANXA2 rendered mice susceptible to chemically-induced carotid arterial thrombosis[90]. To determine the possible involvement of thrombus formation, which is associated with EC injury[91], in cerebral microhemorrhage, we examine all H&E staining and no thrombus was visualized.

We previously showed that the exchange protein directly activated by cAMP (EPAC) plays a critical role during SFGR infections[80] and that EPAC regulates ANXA2-mediated vascular fibrinolysis[92]. Given the regulatory role of EPAC on ANXA2, we retrospectively reviewed archival H&E-stained brain sections of EPAC1-deficient mice[80, 92] infected with an ordinary lethal dose of SFGR, but no histological evidence of CMHs was found, suggesting EPAC is not associated with CMHs during infections.

## Discussion

In this study, we report a novel finding that *ANXA2*-KO mice are more susceptible to rickettsia- or Ebola virus-induced CMHs, underlying mechanisms relevant to endothelial TJs-based BBB dysfunction. Furthermore, we performed a proteomic analysis on the whole brain lysate and isolated endosomes using LC/MS. The obtained list of differentially expressed proteins was subjected to function enrichment analysis, combining GO term and KEGG pathway.

### Whole-brain lysate proteomic functional group

#### Cell-cell junction

BBB properties are primarily determined by BMEC AJs and TJs. Cell-cell AJ junction offers contracting force to bridge adjacent ECs via cadherins and nectins [93]. It physically interacts with TJs and indirectly links to TJ via actin-associated proteins[88, 93]. It is reported that AXNA2 is directly associated with AJ stability. The depletion of ANXA2 dissociates VE-cadherin from AJ[73]. The identified proteins associated with AJs include myosin heavy polypeptide 9 (non-muscle), septins, tyrosine 3-monooxygenase/tryptophan 5-monooxygenase activation protein, HSP90α (cytosolic), aldolase A, enolase 1, heat shock protein 5.These proteins are not direct structural proteins of the AJ such as VE-cadherin and beta-catenin but closely related to AJ assembly and stability. For example, inhibition of HSP90α is capable of attenuate the reduced VE-cadherin and beta catenin level induced by thrombin, LPS, VEGF, or TGF-β1, which all are known to increase endothelial permeability [94–96]. It is also suggested that HSP90α is a downstream effector of RhoA signaling, which is mediated by LPS[96]. Our data shows that HSP90α level was increased 9.4-fold in *ANXA2*-KO brain compared to WT post rickettsia infection. Elevated levels of HSP90α could lead to the instability of the AJ in the presence of LPS[97]. The association of HSP90α and ANXA2 has not been fully understood. There is only one published study supporting the association, showing that high glucose increases *ANXA2* expression in ECs and enhances the association of HSP90α and ANXA2 in both ECs and rat aorta[98]. The functional interaction between ANXA2 and hsp90 in the context of pathogen infection is largely unknown.

#### Stress fiber

Stress fiber is formed by contractile actin filament bundle and a variety of proteins responsible for actin stabilization including septins, α-actin, tropomyosin, Rho-associated protein kinase (ROCK), fascin, and myosin 2, etc.[99]. It is formed in the cells in response to the environmental mechanical stress, therefore only a subset of cells in the body can form stress fiber, such as ECs in the presence of fluid shear stress. Cells cultured on stiff substrate form stress fiber whereas those on the soft substrate have much less fiber. Physiologically, stress fiber is important for cell adhesion, mechanotransduction and AJ maturation[99–101]. Stress fiber plays a positive role in AJ maturation, but it is not always the case in the pathological environment. Such phenomenon is also seen in VEGF- and activated-neutrophil-treated ECs[102, 103].

Our data report four DE proteins in the stress fiber category: septin 5, septin 7, tropomyosin 1 and tropomyosin 3. Three out of the four proteins were increased in *ANXA2*-KO group. The association of ANXA2 to stress fiber formation is elusive. In BHK-IR cells, insulin-mediated loss of stress fiber and resulting cell detachment depend on the presence of AXNA2 and its tyrosine phosphorylation state[74]. Silencing *ANXA2* with siRNA in MOI Muller cells directly leads to the accumulation of stress fiber[70].

### Isolated endosome function group

#### Cell-cell junctions

AJ is a significantly enriched functional group. Ten proteins were found in this category. Endocytosis is known to mediate the stability of AJ[104]. Based on structural analysis using IF and drug intervention, Georgiou et al, suggested that impaired endocytosis leads to AJ instability[104]. AXNA2 is involved in the formation of early endosomes. AXNA2 depletion results in morphological changes in endosomes and alters their distribution[73]. Our proteomic data analyzing the protein difference between WT and *ANXA2*-KO mice upon rickettsial infection identified 10 proteins associated with AJ in the brain endosomal compartment. Interestingly, cadherin complex protein cadherin 2 and talin 1 were upregulated in *ANXA2*-KO after infections. Cadherin 2 is a direct structural protein of AJ, and it is involved in pericyte-EC adhesion, which is critical for the integrity of the BBB. Focal injection of antibody against cadherin-2 is sufficient to induce intracerebral hemorrhage[105]. The biological significance of cadherin-2 upregulation in the endosomal compartment is unknown. It has been shown, however, that cadherin-2 and other several other junction-associated proteins were subjected to rapid endocytosis in a drug-induced disruption of cell-cell contact model [106]. Talin 1 is a well-known AJ-associated protein located at the cytoplasmic side to the AJ[107]. Endosomal talin has been shown to regulate the function of endosomal integrin[108].

#### MHC II protein complex binding and stress response

Three major players are found in MHC II protein complex binding: HSP90α, heat shock protein 8 (HSPa8), and pyruvate kinase. HSP90α is noteworthy. In *ANXA2-*KO mice, it was drastically enriched in whole-brain but decreased in the endosomal compartment, indicating a reduced protein turnover rate or degradation. Given the direct association of ANXA2 and endocytosis, ANXA2 may be an upstream regulator of HSP90α responsible for its recycling upon stimulation. HSP90α is involved in the disruption of BBB function post cerebral ischemic stroke. Inhibition of HSP90α with ATP competitive inhibitor results in reduced activity of metalloproteinase 9 (MMP9), which plays an important role in BBB dysfunction, and rescues TJ protein expression abnormality[109]. HSP90α is also associated with inflammation induced by LPS[110]. It has been shown that the involvement of HSP90α is associated with PI3K and NF-κB pathways[111]. Therefore, a possible mechanism explaining the increased susceptibility of *ANXA2*-KO to *R.australis*-induced CMH is that ANXA2 depletion causes reduced degradation of HSP90α, which destabilizes the cell-cell junction and increases the hyperpermeability of BBB. So far AXNA2 is known to interact with HSP90α and such interaction is increased in the presence of high glucose[98], yet the role of HSP90α-AXNA2 interaction in bacterial infection is unknown. Further *in vitro* experiment is needed to confirm this mechanism.

HSPa8 is a noncanonical member of the heat shock protein 70 family[112]. The majority of HSPa8 resides in the cytosol and nucleus, but upon stimulation, it also translocates to the plasma membrane or exosome, contributing to antigen presentation via MHC II to CD4 T cells, clathrin-mediated vesicles transport, and chaperone-dependent autophagy[112, 113]. In our data, similar to HSP90α, HSPa8 was found to be reduced in the endosomal compartment in *ANXA2*-KO group, whereas it was upregulated in the whole brain lysate (one-fold). However, the biological role of the association between AXNA2 and HSPa8 is unknown.

HSP90α, HSP8 and mitogen-activated protein kinase kinase 4(Map2k4) comprise another functional group, stress response. Map2k4 is a direct activator of c-jun activation protein kinase (JNK) [114]. Map2k4 was enriched in the endosomal compartment in *ANXA2*-KO mice brain compared to WT. Interestingly, it has been reported that *ANXA2* knockdown by shRNA enhances the activation of JNK and p38 in response to oxidative stress[115].

## Conclusion

Our data suggest that 1) ANXA2 per se does not affect the proliferation of *R. australis* or Ebola virus. 2) Also, *ANXA2*-KO does not significantly affect the survival of the mice post-acute *R. australis*. 3) Inflammatory cytokines TNFα and IFNγ are not affected by ANXA2 knock out. 4) ANXA2 is required for stabilizing BBB, especially endothelial TJs during rickettsia and Ebola virus infections. 5) The most relevant functional groups that could be intertwined with the contingent function of ANXA2 are cell-cell junction, stress fiber, and MHC II binding complex, shown by functional enrichment analysis combining the whole-brain lysate and isolated endosome proteome.

Among all the DE proteins we have identified in this study, HSP90α is the most noteworthy because it was present in multiple vascular integrity relevant functional enriched groups. HSP90α is closely related to endothelial hyperpermeability[94]. In our proteomic data, it is upregulated in the whole brain lysate compartment while downregulated in the isolated endosome, indicating that its endocytosis dependent degradation might be impaired.

In conclusion, this is the first report that correlates the special role of ANXA2 and CMHs in the context of rickettsial and Ebola virus infections. Further experiments on the *in vitro* multi-cell co-culture BBB system are necessary to delineate underlying mechanism(s).

## Materials and methods

### Biologic containment

Infectious material and animals were handled in maximum-containment biosafety level 3 (for *R. australis*) and 4 (for Ebola virus) facilities at the Galveston Nationa Laboratory (GNL), University Texas Medical Branch at Galveston.

### Ethics statement

All animal protocols were approved by the Institutional Animal Care and Use Committee of the University of Texas Medical Branch (protocol # 1702018 and protocol # 9505045G). The animal studies were carried out in strict accordance with the recommendations in the Guide for the Care and Use of Laboratory Animals of the National Institutes of Health, USA.

### SFGR mouse infection model[80, 116]

*ANXA2*-KO on C57BL/6 background and C57BL/6 mice were used in this study. Animals (15 WT and 14 *ANXA2*-KO mice) were inoculated with 2LD50 dose (2 × 10^6^ pfu per mouse) of *R. australis* via tail vein injection and observed daily. All procedures followed the approved IACUC protocol. Signs of ruffled fur, hunched posture, labored breathing and closed eyelids were identified as lethal illness (41, 42). The animals were observed for 10 days when most of the animals were all in lethal illness state. For time-dependent pathological study, mice were inoculated with 2 LD50 dose of *R. australis* and euthanized at day 2, 4, 5 post-infection (n=5 for each time point). For mass spectrometry experiments, animals (WT or *ANXA2*-KO) were euthanized at 5 days p.i. and the brain samples were collected and digested into protein lysate for downstream analysis. The time point was selected for LC/MS because this was when mice started to showing up lethal illness.

### Ebola virus mouse infection model[89]

To observe the effect of mice with *ANXA2*-KO compared to WT on Ebola hemorrhagic disease, C57BL6 6-12 week wild type (n=5) and *ANXA2*-KO mice were inoculated intraperitoneally with 50 plaque-forming unit of mouse-adapted strain of Ebola Zaire (Mayinga) virus (provided by Thomas Ksiazek) in 200 uL of PBS. All procedures followed approved IACUC protocol. Mice were monitored multiple times daily for signs of illness and mortality p.i. Daily observations included evaluation of mice for clinical symptoms such as reduced grooming, ruffled fur, hunched posture, subdued response to stimulation, nasal discharge, and bleeding. Tissues and carcasses were collected for downstream assays.

### Sample collection

Blood samples were collected via orbital sinus and serum was obtained after centrifuge and discarding the cellular content of the blood. Complete necropsies were performed on all mice to obtain the organs. For immunohistochemistry or H&E staining, organs were fixed in a 4% (vol/vol) formaldehyde. For mass spectrometry, brain tissues were lysed in protein lysing buffer with proteinase inhibitor and phosphatase inhibitor. Endosome isolation was perform using Minute^TM^ endosome and cell fraction kit.

### IF and H&E staining

The fixed samples were subjected to H&E staining or immunofluorescence (IF) with an antibody against the protein of interest. For IF studies on mouse tissue with mouse antibody, tissue sections were blocked with unconjugated AffiniPure Fab fragment goat anti-mouse IgG (H+L) for 1 hour at room temperature first before incubation with antibodies against ZO-1, occludin, SFG rickettsiae, or Ebola virus overnight at 4 °C, followed by secondary antibody AlexaFluor 594 goat anti-mouse or AlexaFluor 488 goat anti-rabbit antibodies. Before mounting to the cover slide, the tissue sections were stained with DAPI. Normal rabbit and mouse IgGs were used as negative reagent controls (**Supplemental Fig. 1C**). Fluorescent images were taken and analyzed with an Olympus BX51 microscope.

### DNA extraction and rt-qPCR

To quantify the rickettsia loading in the brain tissue, the DNeasy tissue kit (Qiagen, CA, 69506) was used to quantify DNA rickettsial DNA. Briefly, the brain samples were minced into pieces, and subjected to lysis buffer and proteinase k digestion for 10 minutes in 56 degrees Celsius; then DNA was precipitated in ethanol and purified using washing buffer. The purified DNA samples were stored in storage buffer in −20 C°. PCR was performed using the protocol as previously described [117]. Rickettsia-specific citrate synthase (CS) gene (gltA) was used as the target for rickettsia detection (gltA forward: GAGAGAAAATTATATCCAAATGTTGAT; gltA reverse, AGGGTCTTCGTGCATTTCTT)[117].

### ELISA

Plasma samples collected from *ANXA2*-KO and WT mice were used for ELISA to detect TNFα (Qantikine ELISA, MTA00B) and IFNγ (Qantikine ELISA, MIF00). Standard curves were performed using the standard proteins according to the protocol provided in the ELISA kits. The ELISA plates were detected at 450nm.

### LC-MS/MS

Proteins were acetone-precipitated and cleaned with 1 ml of ice-cold wash solution (tri-*n*-butyl phosphate/acetone/methanol (1:12:1 by volume) for 90 minutes and then centrifuged at 2800g for 15 minutes at 4°C. The supernatant was removed and 1 ml ice-cold tri-*n*-butyl phosphate was added and incubated at 4°C for 15 minutes and then centrifuged at 2800g for 15 minutes at 4°C. The supernatant was discarded and 1 ml ice-cold acetone was added and incubated at 4°C for 15 minutes and then centrifuged at 2800g for 15 minutes at 4°C. The supernatant was discarded and 1 ml ice-cold methanol was added and incubated at 4°C for 15 minutes and then centrifuged at 2800g for 15 minutes at 4°C. 50ug of protein was solubilized with 5% SDS, 50 mM TEAB, pH 7.55, in the final volume of 25 uL. The sample was then centrifuged at 17,000g for 10 minutes for debris removal. Proteins were reduced by making the solution 20mM TCEP (Thermo, #77720) and incubated at 65^○^C for 30 minutes. The sample was cooled to room temperature. After 1 uL of 0.5 M iodoacetamide acid was added, the sample was allowed to react for 20 minutes in the dark. 2.75 ul of 12% phosphoric acid was added to the protein solution. 165uL of binding buffer (90% Methanol, 100mM TEAB final; pH 7.1) was then added to the solution. The resulting solution was loaded onto a S-Trap spin column (protifi.com) and passed through the column by a benchtop centrifuge (30-second spin at 4,000g). The spin column was washed with 400uL of binding buffer and centrifuged. The wash was repeated two more times. Trypsin was added to the protein mixture in a ratio of 1:25 in 50mM TEAB, pH=8, and incubated at 37^○^C for 4 hours. Peptides were eluted with 80 uL of 50 mM TEAB, followed by 80 uL of 0.2% formic acid, and finally 80 uL of 50% acetonitrile, 0.2% formic acid. The combined peptide solution was then dried in a speed vac and resuspended in 2% acetonitrile, 0.1% formic acid, 97.9% water and placed in an autosampler vial.

### NanoLC MS/MS Analysis

Instrument performance was verified by analyzing a standard peptide mix and a complex protein digest (HeLa) before the sample set was run between each experimental block and at the end of the experiment. The HeLa data files were analyzed in order to confirm that instrument performance remained consistent throughout the experiment. Peptide mixtures from digested brain tissue were analyzed by nanoflow liquid chromatography-tandem mass spectrometry (nanoLC-MS/MS) using a nano-LC chromatography system (UltiMate 3000 RSLCnano, Dionex), coupled on-line to a Thermo Orbitrap Fusion mass spectrometer (Thermo Fisher Scientific, San Jose, CA) through a nanospray ion source (Thermo Scientific). A trap and elute method was used. The trap column is a C18 PepMap100 (300um X 5mm, 5um particle size) from ThermoScientific. The analytical columns is an Acclaim PepMap 100 (75um X 25 cm) from (Thermo Scientific). After equilibrating the column in 98% solvent A (0.1% formic acid in water) and 2% solvent B (0.1% formic acid in acetonitrile (ACN)), the samples (1 µL in solvent A) were injected onto the trap column and subsequently eluted (400 nL/min) by gradient elution onto the C18 column as follows: isocratic at 2% B, 0-5 minutes; 2% to 32% B, 5-100 minutes; 32% to 50% B, 100-108 minutes; 50% to 90% B, 108-109 minutes; isocratic at 90% B, 109-114 minutes; 90% to 2%, 114-115 minutes; and isocratic at 2% B, till 130 minutes.

All LC-MS/MS data were acquired using XCalibur, version 2.1.0 (Thermo Fisher Scientific) in positive ion mode using a top speed data-dependent acquisition (DDA) method with a 3 sec cycle time. The survey scans (*m/z* 350-1500) were acquired in the Orbitrap at 120,000 resolution (at *m/z* = 400) in profile mode, with a maximum injection time of 50 msec and an AGC target of 400,000 ions. The S-lens RF level was set to 60. Isolation was performed in the quadrupole with a 1.6 Da isolation window, and CID MS/MS acquisition was performed in profile mode using rapid scan rate with detection in the orbitrap (res: 35,000), with the following settings: parent threshold = 5,000; collision energy = 35%; maximum injection time 100 msec; AGC target 500,000 ions. Monoisotopic precursor selection (MIPS) and charge state filtering were on, with charge states 2-6 included. Dynamic exclusion was used to remove selected precursor ions, with a +/- 10 ppm mass tolerance, for 60 sec after acquisition of one MS/MS spectrum.

Database Searching. Tandem mass spectra were extracted and charge state deconvoluted by Proteome Discoverer (Thermo Fisher, version 1.4.1.14). Deisotoping was not performed. All MS/MS spectra were searched using Sequest. Searches were performed with a parent ion tolerance of 5 ppm and a fragment ion tolerance of 0.60 Da. Trypsin was specified as the enzyme, allowing for two missed cleavages. Fixed modification of carbamidomethyl (C) and variable modifications of oxidation (M) and deamidation of asparagine and glutamine, were specified in Sequest. Scaffold (version Scaffold_4.8.7, Proteome Software Inc., Portland, OR) was used to validate MS/MS-based peptide and protein identifications. Peptide identifications were accepted if they could be established at greater than 95.0% probability. Peptide Probabilities from X! Tandem and Sequest were assigned by the Scaffold Local FDR algorithm. Peptide Probabilities were assigned by the Peptide Prophet algorithm [118] with Scaffold delta-mass correction. Protein identifications were accepted if they could be established at greater than 95.0% probability and contained at least 2 identified peptides. Protein probabilities were assigned by the Protein Prophet algorithm [118]. Proteins that contained similar peptides and could not be differentiated based on MS/MS analysis alone were grouped to satisfy the principles of parsimony.

### Functional enrichment analysis

Raw mass spectrometry data was filtered to rule out low abundance protein. Specifically, proteins that are included in the analysis must have at least 4 total spectra count. A list of differentially expressed proteins was obtained by comparing the -spectral counts between *ANXA2-* KO and WT groups and calculating the fold change. Proteins with a fold change greater than or equal to 2 were considered upregulated; protein with fold change less than or equal to −0.6 were considered down-regulated. Selected DE proteins are listed in Table 1-4. Next, the DE proteins were further annotated using The Database for Annotation, Visualization and Integrated Discovery (DAVID) v6.8, which is also capable of doing functional enrichment analysis. Functional enrichment analysis helps us better correlate the molecular patterns with the gross pathology we have observed in the animal model. GO term and KEGG pathways were used to categorize the DE proteins (Figure 1). The Biological Networks Gene Ontology tool (BiNGO) [119] was used to perform the network analysis, which provides a direct structural visualization of the functional enriched groups. Protein-protein interaction (PPI) network was performed using Cytoscape GeneMania[120].

## Acknowledgment

We gratefully acknowledge Dr. Katherin Hajjar for providing the *ANXA2*-KO breeders. We gratefully acknowledge Drs. David Walker and Gerk Volker for reagent support. We gratefully acknowledge Dr. Donald Bouyer for BSL3 facility support. We thank Drs. Shangyi Yu, Xiang Li, Ben Zhang, and Junyin Zheng for technical support. This work was supported by NIH grant R01AI121012 (B.G.) and R21AI137785 (B.G.). The funders have no role in the study design, data collection and analysis, decision to publish, or preparation of the manuscript.

## Supporting Information Legends

Supplemental Figure 1. (A) Comparing the survival of ANXA2-KO (n=15) and WT (n=14) mice challenged by R.australis up to 10 days. No significant difference was found based on Log-rank test. (B) Gross view of the brain surface (R. australis infected WT(top panel) & R.australis infected KO (bottom panel)). Yellow arrows indicate R. australis infected mice brains exhibt a large difference in color.

Supplemental Figure 2. Comparing the survival of ANXA2-KO and WT mice challenged by Ebola virus up to 12 days. No significant difference was found based on Log-rank test, n=5 for both groups, P=0.08.

